# The ribosomal DNA landscape in mammalian muscle during acute and chronic physiological stress

**DOI:** 10.64898/2026.08.13.744441

**Authors:** Daniel Vaughan, Nathanael Wood, Robert Seaborne

## Abstract

Ribosomal DNA (rDNA) is a highly repetitive and complex locus within the mammalian genome that exhibits substantial inter-individual variation in number of rDNA copies and epigenetic regulation. Nonetheless, our understanding of rDNA biology in skeletal muscle during periods of physiological stress is limited. Using publicly available whole genome and reduced representative bisulfite sequencing data sets, we identify a concurrent reduction in both the number of rDNA copies and the methylation profile of the rDNA in aged vs young mice, supported by large effect sizes and permutation testing, with significant reductions in methylation of the 18S coding unit in aged, compared to young controls (*p* = 0.024). We found a strong positive correlation between rDNA copy number and 18S methylation across both young and aged mice (*p* = 0.004; Spearman rho = 0.842). After analysing publicly available muscle (skeletal and cardiac) data sets following acute insult (endurance exercise, cancer cachexia, spinal cord injury), we do not observe a similarly coordinated epi-genetic modification in rDNA biology but uncover tissue and sex-specific differences in rDNA copy number or methylation status, in isolation. These findings suggest ageing as a unique physiological insult in which coordinated epi-genomic remodelling of the rDNA region appears, representing a previously underappreciated feature of the muscle ageing trajectory.

## Introduction

The maintenance in integrity, function and structure of skeletal muscle is of paramount importance for overall organism health- and lifespan. Skeletal muscle is also remarkably malleable. Indeed, the tissue shows significant plasticity, (mal)adapting at molecular, morphological and functional levels, to array of physiological stressors (Smith et al., 2023). Most prominent of these is the associated degradation in function, mass and integrity of muscle during the ageing process (Sayer et al., 2024). Exactly what governs this adaptative capacity remains incompletely understood.

There is considerable interest in the role of the ribosome in orchestrating adaptation to physiological stress, for which, skeletal muscle ribosome biology has received limited attention. The ribosome consists of a large (60S) and small (40S) ribonucleoprotein unit, collectively maturing into the 80S functional ribosomal complex (Rodriguez-Algarra et al., 2025). This complex consists of ∼80 ribosomal proteins (RPs) and four nascent ribosomal RNA (rRNA) transcripts, three of which (18S, 5.8S and 28S) are coded from the 45kb ribosomal DNA (rDNA) locus on the p-arms of acrocentric chromosomes (e.g. Chr:12, 15, 16, 18 and 19 in *Mus musculus*; Figure 1A), that exist in multiple copies (Rodriguez-Algarra et al., 2025). The multi-copy, highly repetitive nature of rDNA sees it typically excluded from traditional genomic analyses, nonetheless recent growing evidence suggests that the rDNA locus is a site of remarkable heterogeneity and variability (Rodriguez-Algarra et al., 2022), giving rise to functional and physiologically important outcomes. For example, we have previously shown that fundamental rDNA copy number associates with post-pubertal rodent growth rate and with body mass index in humans (Law et al., 2024). The rDNA locus is also epigenetically sensitive (Srivastava et al., 2016a), with DNA and chromatin marks rendering transcriptionally active, silent or ‘poised’ rDNA copies. Given skeletal muscle is the most adaptive tissue in the mammalian organism, it is plausible that the epigenetic and genetic status of the rDNA locus change following physiological stress.

**Figure 1.**
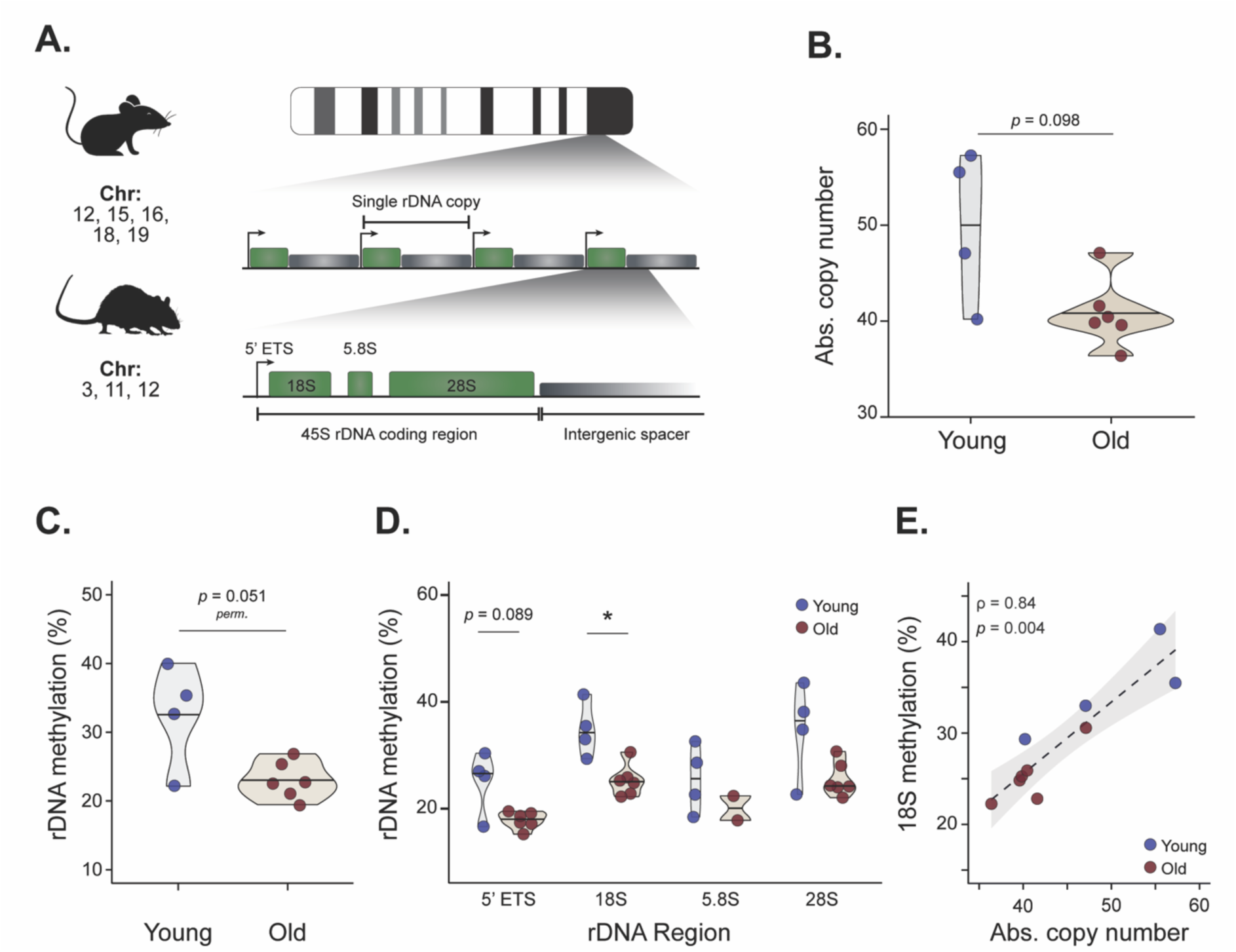
**A.** Representative schematic of mouse and rat rDNA genomic location, its palindromic repeat architecture and the coding regions within each rDNA copy. **B**. Absolute rDNA copy number of young and old mice calculated from WGBS data. **C**. The 45S DNA methylation of young and old mouse skeletal muscle and the methylation status of the 5’ETS and rRNA coding units of the rDNA. **D.** A significant association between rDNA copy number and DNA methylation of the 18S unit across both young and old mice. Statistical values are presented for N=4 young and N=6 old mice, except for the 5.8S methylation data (D; N=2 old). **E.** A positive correlation between absolute rDNA copy number and 18S methylation and positive correlation from old to young.

Nonetheless our understanding of rDNA dynamics during physiological stress, such as the impact of ageing, remain limited. Previous work in young, aged and aged + exercise in mice suggest a complex epigenetic story, with CpG specific methylation either increasing or decreasing in methylation (hyper/hypo-methylation, respectively), but a more pervasive hypermethylated response (Murach et al., 2022). Interestingly, this work identified a small sub-set of differentially methylated CpG sites that, following exercise in the aged mice, reversed their epigenetic signature, a finding our lab has also observed in multi-copy genomic regions (Ruple et al., 2021). However, the highly repetitive nature of such loci makes traditional differentially methylated analyses complicated at CpG resolution and no global/coding unit level analyses was previously conducted. Furthermore, no known work has identified rDNA copy number dynamics in ageing skeletal muscle. Analyses of the rDNA region from an epigenetic and genetic perspective, following acute exercise reveals a further complexity in understanding the role that rDNA plays in supporting muscle adaptation (Figueiredo et al., 2021; Godwin et al., 2024)

Research into the functional role and epi-genomic behaviour of the rDNA locus in the post-mitotic terminally differentiated muscle tissue, while in its infancy, is complex. Here, using publicly available whole genome and reduced representative bisulfite sequencing (WGBS and RRBS, respectively) data sets across rodent models, we first analysed the rDNA epi-genomic dynamics in aged skeletal muscle revealing tandem modifications at both genomic and epigenomic levels. We further explored both negative and positive physiological stressors, in skeletal and cardiac muscle, to understand how the rDNA locus behaves following relatively acute insult.

## Results

### Ageing in mouse skeletal muscle is associated with a reduction in absolute rDNA copy number and methylation status

We have previously used whole genome / reduced representative bisulfite sequencing data (WGBS/RRBS respectively) to identify an association between absolute rDNA copy number and physiological outcomes (Law et al., 2024). Here, using publicly available data (Oyabu et al., 2025), we report a trend for reduction in relative rDNA CN (Fig. 1A) in skeletal muscle of old mice (C57BL/6NRcl; N=6; 40.8 ± 3.5) compared to young (N=4; 50.0 ± 7.9) (*p*=0.098; Fig. 1B). While not reaching statistical significance, a large effect size between ages was identified (Hedges’ g = 1.27), suggesting a meaningful reduction in absolute rDNA CN.

Using the associated WGBS data at 50X coverage, we next investigated whether ageing in mouse skeletal muscle modified the epigenetic landscape of the rDNA region (Fig 1A). The entire coding unit showed a reduction in DNA methylation consequential to ageing in mouse skeletal muscle (*p* = 0.082; permutation testing *p* = 0.051; Fig. 1C). Closer inspection of the data revealed the strongest reduction in methylation from young to old skeletal muscle in the 18S coding region (*p* = 0.024) and in the 5-prime externally transcribed spacer (5’ETS, *p* = 0.082; Fig. 1D). Absolute rDNA copy number was significantly associated with the methylation status of the 18S region in skeletal muscle across both young and old through multiple statistical tests (Spearman rho = 0.842, *p* = 0.004; Pearson r = 0.908, *p* = 0.0002; Fig. 1E), with rDNA copy number explaining 82% of the variance in DNA methylation (R^2^ = 0.824).

### Epigenetic, but not genomic, changes in rDNA landscape following exercise in striated muscle

Ageing skeletal muscle represents a chronic stress exerted on the tissue. After identifying trends for reduction in both copy number and methylation profile of the rDNA locus, we sought to examine how an array of acute physiological stressors modifies the rDNA landscape in muscle.

To do so, we downloaded gastrocnemius and cardiac RRBS data from the MoTrPAC study (Sanford et al., 2020), containing control rats (Fischer 344) and rats exercised across 1, 2, 4 and 8 weeks with N=5 male and N=5 female across all groups (MoTrPAC Study Group et al., 2024). A two-way ANOVA by tissue identified no significant effect of exercise training time-point in either cardiac (*p* = 0.101) or gastrocnemius (*p* = 0.476) tissue. However, we did find a significant effect of sex on rDNA copy number in cardiac tissue (*p* < 0.0001; Fig. 2A), but not in gastrocnemius skeletal muscle (*p* = 0.101). There was no interaction of sex by exercise in either tissue (cardiac, *p* = 0.845; gastrocnemius, *p* = 0.568).

**Figure 2.**
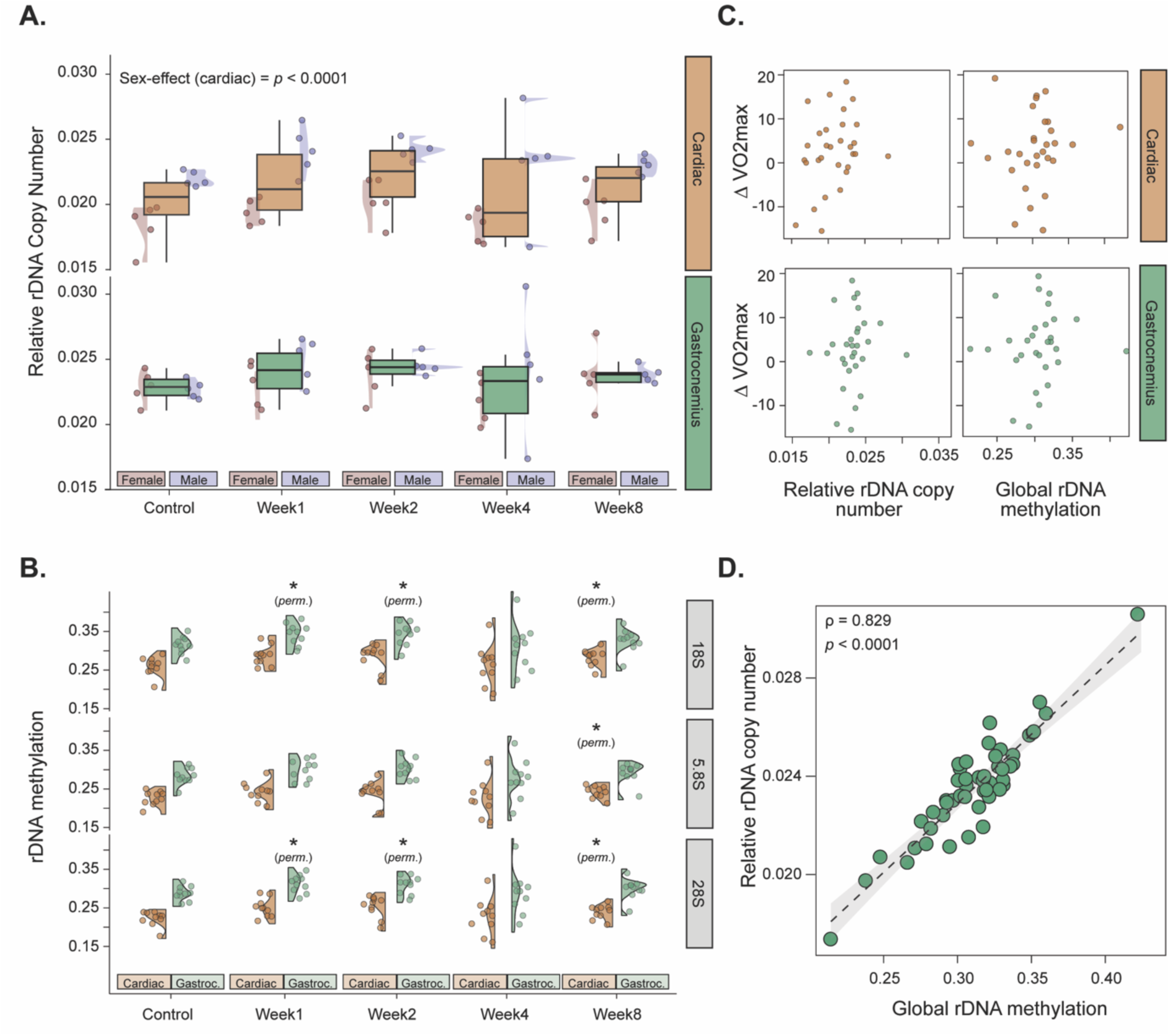
rDNA regulation in cardiac and skeletal muscle from the MoTrPAC 6-month-old rat endurance training programme. **A.** Relative rDNA copy number for cardiac and gastrocnemius tissue across control and 8-weeks of training. **B.** rDNA methylation in three coding units (18S, 5.8S, 28S) of the rat genome, in both cardiac and gastrocnemius rat tissue. Permutation tests (see methodology) show significance against tissue-specific controls. **C.** Associations of changes in relative rDNA copy number (left panel) and global rDNA methylation (right panel) for cardiac (top panel) and gastrocnemius (bottom panel) against changes in ΔVO2max of rats compared to control. **D.** Significant correlation between relative rDNA copy number and global rDNA methylation in rat skeletal muscle, only. Statistical values are presented for N=5 male and N=5 female for all time points for cardiac and gastrocnemius tissue, except Figure C. in which only control, week 4 and week 8 data are presented.

The current *Rattus norvegicus* assembly contains the coding units of the rDNA locus only, that is the 18S, 5.8S and 28S (Fig 1A). We inspected the methylation profile of both skeletal and cardiac tissue of these coding regions across the exercise time-course within the MoTrPAC study. Combining sexes, we observed a modest increase in global rDNA methylation in skeletal and cardiac tissue, Kruskal effect size analyses (ε²=0.104 and ε²=0.121, respectively) suggesting potentially meaningful biological differences. Permutation testing identified weeks 1 and 2 of endurance training (vs control) to have the biggest impact on rDNA methylation, with gastrocnemius tissue showing an increase in both the 18S (*p* = 0.0179, t = −2.5746; *p* = 0.0413, t = −2.2474, week 1 and 2 respectively) and 28S (*p* = 0.0128, t = −2.7136; *p* = 0.0214, t = −2.5485, week 1 and 2 respectively; Fig. 2B). Whereas in the cardiac tissue, the 28S was significantly increased in weeks 1 and 2 (*p* = 0.0156, t = −2.64; *p* = 0.0283, t = −2.3768, respectively; Fig. 2B). Interestingly, in cardiac tissue, 8 weeks of endurance training increased rDNA methylation in all three coding units (18S, *p* = 0.0306, t = −2.3093; 5.8S, *p* = 0.0327, t = −2.2816; 28S, *p* = 0.0313, t = −2.2574; Fig. 2B). Collectively, our analyses suggest a potential increase in rDNA methylation in the acute period (week 1 and 2) following endurance exercise in gastrocnemius and cardiac striated muscle.

We correlated changes in rDNA biology with changes in VO2max (pre to post, Δ), as one of the most pertinent physiological measures associated with endurance performance/adaptation. After downloading phenotype data from the MoTrPAC consortium, only control week 4 and week 8 data were available for VO2max. Subsequently, neither rDNA methylation nor relative rDNA copy number were statistically associated with changes in VO2max (Fig. 2C). Similarly to ageing skeletal muscle (Fig 1E.), a significant correlation was observed between methylation and copy number of skeletal muscle rDNA (Spearman ρ=0.829, *p* < 0.0001; Pearson r = 0.917, *p* < 0.0001; Fig. 2D). Interestingly, no rDNA methylation and copy number association was found in cardiac tissue (Pearson r = 0.026).

### Progressive rDNA copy number, but heterogeneous methylation changes in cachexic skeletal muscle

We last wanted to observe the rDNA landscape of skeletal muscle in the context of cancer associated muscle catabolism (cancer cachexia). Cancer notoriously induces large effects on the ribosomal DNA landscape of affected micro-environments and cells (Xu et al., 2017). Less is known about tissues who succumb to secondary effects of cancer, such as skeletal muscle that undergoes significant deterioration of both mass and functionality. We downloaded a cancer cachexia mouse RRBS data set (Cabrera et al., 2026), where male (N=8) and female (N=8) mice (BALB/c) were injected with Colon-26 (C26) cell allografts for a period of 10, 20 and 25 days of analyses, compared to PBS injected controls (N=16 samples/condition). Compared to control, we observed a progressive increase in relative rDNA copy number in 10, 20 and 25 days cachexic skeletal muscle (25day vs control, *p* = 0.051) which, following permutation testing, reached significance at 25days (*p* = 0.002). Post-hoc analyses revealed significance was attained in female (Benjamini-Hochberg, *p_ad_*_j_= 0.039) but not male mice (Benjamini-Hochberg, *p_ad_*_j_= 0.667). Both 10days and 20days of cancer did not statistically alter rDNA copy number compared to control.

At a methylation level, we observed very little difference by cachexia time-point, compared to control across the rDNA unit (45S; Figure 3B;) or in specific rDNA regions (supp file X). However, we found that control males had statistically higher rDNA methylation (*p* = 0.045) across the rDNA unit compared to female control mice (Figure 3B). This sex-specific difference is not maintained in any time-point of cancer cachexia (Figure 3B). Fligner-Killeen homogeneity tests revealed a significant difference in the profiles and uniformity of rDNA methylation in the 18S (χ² = 12.12, *p* = 0.006995) indicating unequal variance in this region, across the cachexic groups. No other statistically significant observations in rDNA methylation homogeneity were observed. No correlation was identified between copy number and methylation across the rDNA when analysing by condition, sex or rDNA region (Figure 3C).

**Figure 3.**
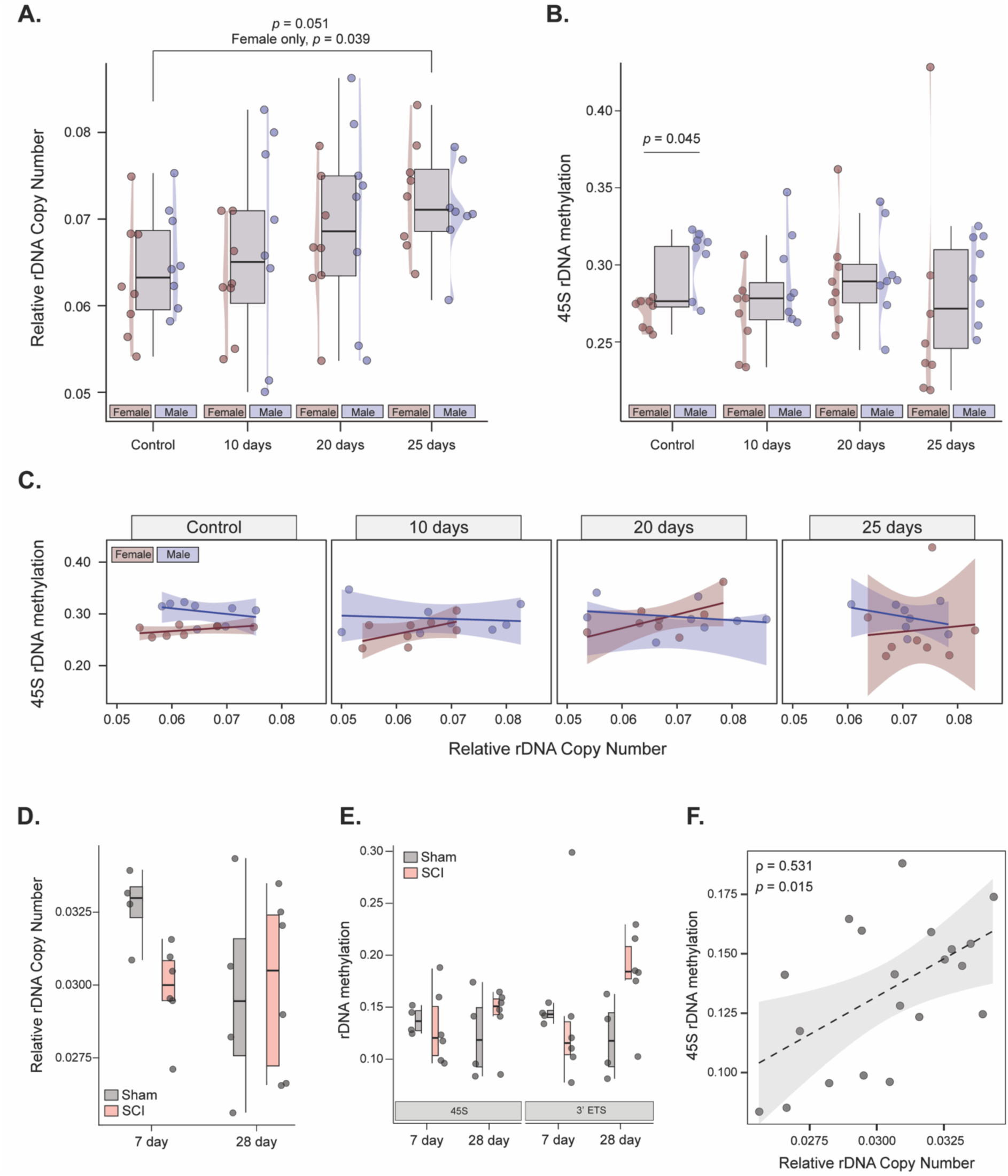
Relative rDNA copy number, DNA methylation and their association in catabolic skeletal muscle. **A.** Relative rDNA copy number and 45S rDNA methylation (B) in mice of control (N=8 male, N=8 female) and following 10, 20 and 25 days of C26 cell treatment, with no significant associations observed between 45S rDNA methylation and relative rDNA copy number in any condition (C). **D.** Copy number of male sham control (N=4) and 7-day (left) and 28-day (right) spinal cord injury (N=6). **E.** rDNA methylation profiles across the 45S (left) and the 3’ETS (right) in sham and cachexic skeletal after 7 and 28-days of treatment. **F.** A significant positive association observed between rDNA methylation and relative rDNA copy number in all mice. Significance values represented in figure panel. Statistical analyses, see methodology.

### Skeletal muscle rDNA methylation remains stable during acute and chronic periods of spinal cord injury muscular catabolism

We last obtained spinal cord injury and sham control mouse skeletal muscle RRBS data (C57BL/6NRcl), across 7-days and 28-day worth of insult (Potter et al., 2023), to observe the dynamics of rDNA during progressive skeletal muscle catabolism separate from cancer (Fig 3A-C). Here, we identified relatively stable rDNA genetic and epigenetic behaviour (Fig 3D-E). Indeed, statistical analyses revealed little relative rDNA copy number alterations from sham control to SCI in either time-points (Fig 3D) and while effect size analyses and permutation testing identified relative rDNA copy number increase in 7-day SCI vs 7-day sham control (*p* = 0.0133, t = −3.11) we believe this to be driven by artefacts in the control condition (7-day sham) only. Indeed, moderate-large effects were also observed when comparing sham controls of 7-day and 28-day (*d* = −1.07), with no observed effect between 7-day SCI and 28-day sham control (*d* = −0.05). At a methylation level, permutation testing revealed a small reduction in rDNA methylation within the 3’ETS region (*p* = 0.046, t = 2.39) after 28days of SCI (vs relevant sham), which failed to reach significance following multiple testing (Benjamini-Hochberg, *p*adj = 0.35). No other significant findings were identified via permutation testing, nor acknowledged via effects size analyses. Similarly to previous skeletal muscle data sets, we did identify a positive association between rDNA methylation across the 45S unit and relative rDNA copy number in all data points combined (*p* = 0.015, Spearman ρ = 0.531). Collectively, these data suggest that mouse skeletal muscle rDNA remains relatively stable during both acute and chronic periods of SCI.

## Discussion

Using publicly available data sets, we explored the dynamic genetic and epigenetic response of ribosomal DNA in mammalian muscle during periods of physiological perturbation. Skeletal muscle ageing was uniquely associated with coordinated reductions in both rDNA copy number and DNA methylation across the coding unit, maintaining a positive correlation between genetic and epigenetic status of the tissue. This shared genetic and epigenetic response is not found in any other stress insult. Indeed, our analyses show that endurance exercise training induces increased DNA methylation of the skeletal muscle rDNA unit but no changes in relative rDNA copy number changes and skeletal muscle of cancerous mice display increased relative rDNA copy number without changes in rDNA methylation. Collectively, our findings suggest that rDNA is not uniformly responsive to physiological stress but undergoes distinct genetic and epigenetic remodelling dependent on the intensity and longevity of insult. In ageing, skeletal muscle absolute rDNA copy loss and hypomethylation may represent a compensatory mechanism that preserves habitual transcriptional capacity, but at the expense of rDNA genomic buffering capacity

Ageing is accompanied with genome-wide epigenetic modifications in skeletal muscle that are purported to underpin some of the age-associated loss in skeletal muscle function and health (Sharples et al., 2018). Traditional analyses of methylome wide data sets focus on the coding regions of the genome, with multi-copy, repeat regions typically excluded from analyses. As such, our understanding of the genetic-epigenetic relationship of these regions, in muscle, during periods of physiological insult, is sparse. Here, we analysed a publicly available post-bisulfite adapter tagging (PBAT) mouse aged data set (Oyabu et al., 2025) alongside computational pipelines to assess both absolute rDNA copy number and rDNA methylation status, as per our (Law et al., 2024; Rodriguez-Algarra et al., 2022) and previous labs have done (Gibbons et al., 2015). Firstly, we identified a trend in reduction for absolute rDNA copy number in aged compared to young mice, which to our knowledge, is the first work to identify reduced absolute rDNA copy number with ageing in skeletal muscle, which is particularly interesting given the post-mitotic, terminally differentiated and multi-nucleated characteristics of skeletal muscle cells (Frontera and Ochala, 2015). It is believed that most incidence of rDNA copy number loss or shrinkage occur during unequal homologous recombination and excision (Nelson et al., 2019), a process typically conserved for mitotic DNA replication in somatic cells. Given the post-mitotic nature of skeletal muscle cells, an alternative mechanism is likely to underpin rDNA copy number loss. Two mechanisms are transcription-associated recombination and excision, and transcriptionally derived RNA:DNA loops (R-loops), both of which occur in contexts of intense rDNA transcription. Given the high translational demands of skeletal muscle, it is conceivable that sustained rDNA transcription could contribute to age-associated rDNA instability through transcription-associated recombination or R-loop formation. Whether such mechanisms contribute to rDNA loss in post-mitotic skeletal muscle remains unknown, and future work is needed to explore these potential mechanisms.

It is now well established that mammals show vast inter-individual variation in rDNA copy number (Gibbons et al., 2014; Parks et al., 2018), with the physiological role of this variety, yet not fully determined. It is also well established that, while rDNA copy number may vary, only a relatively small number of these copies are under active transcription. Indeed, rDNA is known to be regulated in a mosaic pattern of several epigenetic modifications (including DNA methylation and at the chromatin level) that render rDNA units in either silenced, active or in a ‘poised’ state of transcriptional behaviour (Potapova et al., 2025; Srivastava et al., 2016b), where hypomethylation of rDNA is associated with transcriptionally active rDNA units (and vice versa). Here, using the same PBAT data, we report a concomitant reduction in rDNA methylation and absolute rDNA copy number reduction in aged mice, so much so that a maintained positive association between copy number and methylation was identified across young and aged samples. Previous work has shown that ageing is associated with pervasive hypermethylation, with rDNA epigenetic clocks being a robust mechanism for epigenetic ageing analyses (Wang and Lemos, 2019). Our findings in skeletal muscle are contradictory to this, suggesting the rDNA epigenetic-genetic dynamics in skeletal muscle tissue behave differently to that of traditional tissues. The only known previous work examining rDNA methylation in skeletal muscle reports both hyper- and hypo-methylated CpG methylation response in aged mice (Murach et al., 2022). Differences in methylome data sets PBAT vs RRBS (Olova et al., 2018) and in the analyses of these data (CpG vs region specific methylation) may be responsible for the differences observed in these analyses. Nonetheless, our findings pose several intriguing possibilities, for which we propose a model in which rDNA copy number reduction is an active outcome of the ageing process, reductions in DNA methylation may help preserve the transcriptional outputs of the remaining rDNA array. The overall rDNA transcriptional behaviour of aged mice may therefore be unaltered due to this epigenetic-genetic trade-off, but the number of reserve rDNA copies that can become activated if required, are significantly reduced. While plausible, these hypothetic remarks must be considered with caution as our analyses does not directly analyse active or inactive rDNA copies, nor in the response of aged mice to physiological stress. Future work using both chromatic and DNA level epigenetic assays to understand the rDNA epigenetic-genetic behaviour in ageing, is warranted.

After identifying a tandem relationship between absolute rDNA copy number and methylation status in aged skeletal muscle, we sought to determine whether a similar relationship exists in muscle during periods of relatively acute stress, compiling an array of data sets to examine the response. Using the MoTrPAC rat endurance exercise data set and following permutation testing, we identified an increase in rDNA methylation without change in copy number, albeit we did identify a strong different in rDNA copy number between gastrocnemius and heart samples of the same rats (Figure 2). Previous work from the von Walden lab showed that, following endurance exercise in mice 45S pre-rRNA levels were significantly reduced compared to control, while resistance exercise had a time-dependent increase in the rRNA transcript expression (Figueiredo et al., 2021). These authors used an array-based method to inspect promotor methylation levels, identifying a time-dependent increase in promotion rDNA methylation. Our analyses support and extend these findings, identifying increase rDNA methylation across the coding units of rats during prolonged endurance exercise training, which, interestingly does not associate with changes in VO2max at 4 or 8 weeks. At a genomic level, we identified a clear copy number difference between cardiac and gastrocnemius tissues in the MoTrPAC data set (figure 2), with only gastrocnemius muscle showing a strong positive correlation between rDNA methylation and relative rDNA copy number. While this might be due to the highly heterogenous cell composition of the heart tissue confounding our analyses, as we have previously discussed (Seaborne and Ochala, 2023) there may also be a fundamental biological rationale for this. Skeletal muscle is remarkably and essentially plastic, continually adapting to differing stimuli (workload, mechanical stress, nutrient availability) and its ability to adapt relies on synthetically meeting the demands of the milieu, at a protein level (Egan and Zierath, 2013). The absence of this relationship in cardiac tissue may reflect fundamental differences in rDNA regulation between tissues, although differences in cellular composition and tissue architecture represent important alternative explanations. Again, future work is needed to understand the epigenetic-genetic dynamics in cardiac tissue. Our analyses also show no discernible correlation between relative rDNA copy number in either gastrocnemius or cardiac tissue, with VO2max as a key physiological marker of endurance performance and training adaptation (Hawley et al., 2014). Nonetheless, this is the first time this association has been explored in endurance exercise, with previous work finding a similar lack of association in the anabolic response following hypertrophic stimuli in humans (Godwin et al., 2024). Superficially these data would suggest that rDNA copy number may not have an active role in exercise adaptation of skeletal muscle. However, no work has yet determined whether the number of transcriptionally active rDNA copies is the key adaptive marker of skeletal muscle following exercise, which requires future work. Collectively, we show tissue specific epigenetic changes following exercise without changes in fundamental rDNA copy number, suggesting that chronic, life-long associated physiological stress is necessary to inflict tandem epigenetic-genetic changes in muscle rDNA biology.

Similarly to our endurance exercise analyses and unlike the ageing data, we did not find associative changes in relative rDNA copy number and methylation status in either the skeletal muscle cancer (Cabrera et al., 2026) or spinal cord injury analyses (Potter et al., 2023), both reflecting catabolic insults. Interestingly, however, we do identify a progressive increase in relative rDNA copy number in cachexic skeletal muscle (Figure 3). Plethora of previous work has identified that the rDNA locus becomes incredibly unstable during varying types of cancer where, commonly, both losses and gains of rDNA copies (Smirnov et al., 2021) and heterogenous and complex epigenetic behaviour is observed. Our findings partially support this. The observation that rDNA copies increase in skeletal muscle of mice injected with C26 cancer cell allografts is intriguing. In this work, skeletal muscle is not the primary site of disease, but is a secondary catabolic site reflecting the cachexic behaviour of the tissue during cancer. Thus, while previous work shows both loss and gain of rDNA copies in cancerous cells (Wang and Lemos, 2017) our analyses reflect the rDNA behaviour of a secondarily diseased catabolic tissue. We do find an increased heterogeneity of rDNA methylation in the 18S of cachexic skeletal muscle (compared to control), that likely reflects the complex nature of cancer in rDNA biology.

Our analyses of publicly available data, suggest tandem epi-genetic modification in aged mouse skeletal muscle, which is not observed in more acute physiological insults. Our observations, while interesting, require caution. First, all data sets herein are derived from part/whole muscle tissue samples, capturing the composite of tissue cell types and muscle cell sub-types (Seaborne and Ochala, 2023). It is difficult to delineate the precise contribution of different cell types/sub-types within each biopsy and condition analysed, and, given that the cellular composition of these changes from condition-to-condition (Seaborne and Ochala, 2023), it plausible that rDNA epi-genetic observations here are not reflective of muscle cell biology. Indeed, recent work from our group has shown distinct differences in single myofibre rDNA biology at the proteomic level (Moreno-Justicia et al., 2025). Thus, future work must use single myofibre-based approaches to inspect rDNA biology across the ageing trajectory and following acute insult. Throughout the analysis of the various datasets included in this study, we have utilised permutation testing to ascertain whether there is significance across conditions and subsequent rDNA genomic regions. Permutation testing serves as a non-parametric technique for exact inference of p-values for a given test statistic, enabling us to reject the null hypothesis (Huang et al., 2006). Permutation tests are used in clinical trials to reject the null hypothesis for treatment efficacy and can aid with the obstacle of small sample sizes (Unseld et al., 2025), however, there are limitations. Permutation testing is known to have instability in p-values, an inability to adjust for covariates, and is conservative, failing to reach small p-values given the discrete nature (Berger, 2000). Future work should therefore increase sample sizes to validate the findings and outcomes observed in our work. The resolved copy number estimating in our ageing analysis is lower than previously reported in mouse data (e.g. (Rodriguez-Algarra et al., 2022), which may, in-part be due to fundamental differences in sequencing library preparations. Indeed, the ageing data set (Oyabu et al., 2025) used a post-bisulfite adaptor tagging (PBAT) WGBS library preparations (Miura et al., 2012), generating alignment rates of ∼43%, in our hands. The remaining data sets in our analyses use the RRBS approach (Meissner, 2005) and, following our analyses, have a higher alignment rate (ranging from ∼60-70%). Thus, future work in ageing skeletal muscle should use more contemporary DNA methylome based pipelines to ensure alignment and coverage do not impede analyses and data interpretation.

Collectively, our analyses of publicly available data sets highlight ageing as a chronic physiological stress leading to a reduction in both rDNA copy number and rDNA methylation, that may reflect a maintenance of active rDNA copies but a reduction in rDNA buffering capacity. No acute physiological stress induces the same extent of genetic-epigenetic modifications in muscle rDNA. Future work using chromatin and long-read sequencing approaches/assays, should now expand these findings to inspect active, poised or silenced rDNA copies and their role in skeletal muscle ageing biology.

## Methodology

### Data sets

All data (whole genome- and reduced representative bisulfite sequencing) analysed in this work was publicly available, downloaded from The Gene Expression Omnibus (GEO). The MoTrPAC physiological/phenotype data was downloaded directly from the consortiums github: https://github.com/MoTrPAC. Data sets used for this work are listed table 1.

**Table 1.** Publicly available data sets used in this study.

| Experiment | Data type | Species | Strain | Reference | Notes |
| --- | --- | --- | --- | --- | --- |
| Skeletal muscle ageing | WGBS | Mus musculus | C57BL/6 NRcl | (Oyabu et al., 2025) | Post-bisulfite adapter tagging |
| Skeletal muscle endurance exercise | RRBS | Rattus norvegicus | Fischer 344 | (MoTr et al., 2024; Sanford et al., 2020) | - |
| Cardiac/heart endurance exercise | RRBS | Rattus norvegicus | Fischer 344 | (MoTr et al., 2024; Sanford et al., 2020) | - |
| Skeletal muscle cancer cachexia | RRBS | Mus musculus | BALB/c | (Cabrera et al., 2026) | C26 arm of experiments analysed |
| Skeletal muscle atrophy | RRBS | Mus musculus | C57BL/6 NRcl | (Potter et al., 2023) | Control and spinal cord injury conditions analysed |

### Genome preparation

The reference ribosomal DNA build is a single linear sequence, unlike the in-situ repeat palindrome, complicating short-read alignment. To enable alignment of reads across the 5’ ETS and to avoid mis-/ mulit-alignment of rDNA reads, and rDNA looped reference was generated, and competitive alignment loci were masked, similarly to our recent work (Law et al., 2024; Rodriguez-Algarra et al., 2022). First, in the *Mus musculus* genome, the rDNA unit was looped in silico by rotating the sequence at a defined ‘breakpoint’ located in the IGS sequence (position 35000), permitting a contiguous alignment across the repeat junction. We need removed and masked spurious alignment loci that shared highly similar matches with the canonical rDNA sequence (using BLAST [blastn, task = megablast], percent identify of ≥ 97 %, alignment length >249bp, E-value threshold 1e-20). Hits were converted BED formant and merged using bedtools merge. This masking strategy was designed to eliminate genomic loci with sufficient homology to attract rDNA-derived reads, while minimizing unnecessary masking of divergent or low-complexity regions. For *Rattus norvegicus*, an identical process was followed with the exception that no rDNA looping was performed due to the constrained rDNA reference, containing only the 18, 5.8 and 28S without flanking and intergenic sequences. The final reference genomes were generated by concatenating the masked assembly (with the looped rDNA contig for mouse) and indexed (samtools faidx) ready for short read alignment.

### Data processing

Raw FASTQ files were assessed for quality using FastQC (v0.11.9). Adapter and low-quality base trimming was performed with Trim Galore! (v0.6.6) using the options −-rrbs, −- non_directional (where applicable), and −-paired (where applicable) for bisulfite sequencing. Trimmed reads were aligned to the *Rattus norvegicus* rn7 or *Mus musculus* mm39 reference genomes using Bismark (v0.23.0) with Bowtie2 (v2.5.1) as the underlying aligner. Alignments were processed with SAMtools (v1.17) for sorting and indexing. PCR duplicates were removed using deduplicate_bismark, and methylation calls were extracted with Bismark’s methylation extractor (options: −-non_directional (where applicable), −-bowtie2, −- comprehensive, −-bedGraph, −-counts). All analyses were executed in a modular environment managed by Python (v3.11.6).

### Absolute and relative rDNA copy number

For all datasets included in this study the relative rDNA copy number was calculated and for the **<u>aged</u>** dataset the absolute rDNA copy number was calculated given this included WGBS methylation data. The relative rDNA copy number (*R*) is calculated using the following formula,

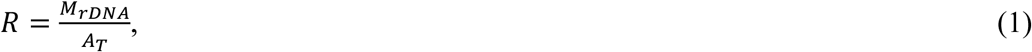

where *M_rDNA_* is the number of mapped reads for rDNA post alignment and *A_T_* is the total number of aligned reads. These numbers can be collected from the Bismark alignment and the SAMtools idxstats report respectively. Similarly, these values can be determined from the coverage values extracted in methylKit (Akalin et al., 2012), as demonstrated in the package myoRDNA. The absolute rDNA copy number (*A*) is calculated using,

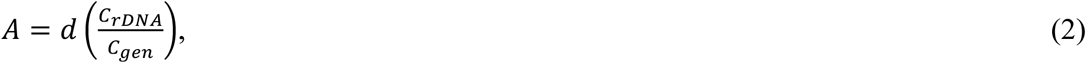

where *d*is the factor depending on if the sample is haploid (*d* = 1) or diploid (*d* = 2), *C_rDNA_* is the summation of the coverage across all positions located in the rDNA chromosome and *C*_g*en*_ is the summation of the coverage across all positions in the non-mitochondrial chromosomes. The code for determining the absolute rDNA copy number is part of the Mordan repository.

### rDNA methylation values

Across all datasets and samples, the methylation values are calculated using the following formula,

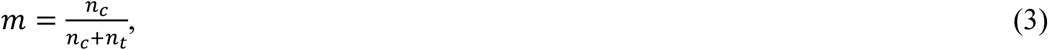

where *n_c_* is the number of methylated Cytosines at a position and *n_t_* is the number of Thymines at a position. The values for *n_c_* and *n_t_* are obtained through the methylKit package. *m* is subsequently averaged over all positions for a specific rDNA region of interest. The regions for averaging across the rDNA contig using genomic positions in mouse are 45S:1-13,403, 5’ETS: 1-4,007, 18S: 4,008-5,877, 5.8S: 6,878-7,034, 28S: 8,123-12,852, 3’ETS: 12,852-13,403, ITS-1: 5,878-6,877, and ITS-2: 7,035-8,122. For the rat genomic ranges are 18S: 1-1,872, 5.8S: 1,873-2,029, and 28S: 2,030-6,836.

### Statistical analysis

Methylation levels across rDNA regions were analyzed in R (v4.5.3). Raw methylation counts were processed using methylKit (v1.36.0, Bioconductor v3.22) to calculate methylation proportions (beta values) for each CpG site. Aggregated methylation levels for predefined rDNA regions were computed, and statistical comparisons between groups (e.g., control vs treatment) were performed using rstatix (v1.0.0). Each dataset was assessed for normality through the Shapiro—Wilk test, and homogeneity of variance was determined using the Fligner—Killeen and Levene’s test. Non-parametric tests, (Wilcoxon rank-sum test and Kruskal-Wallis test), were applied to assess differences in methylation levels, accounting for the non-normal distribution of the data. This was performed in conjunction with one-way ANOVAs for normal and homogeneous variances and Welch’s t-test otherwise. The effect size was analysed across conditions to observe variation in the datasets using the eta-squared metric for the Kruskal-Wallis test and Cohen’s D for the ANOVA test. A two-way ANOVA (sex and time-point) was used to assess relative rDNA copy number across MOTR-PAC tissue data sets, and across cachexic rDNA methylation data. Post-hoc analysis performed using the Tukey HSD or Games-Howell test with Benjamini-Hochberg adjustment. MKInfer (v1.3) was used specifically for permutation testing to further validate the robustness of the observed differences, all permutation tests were two-sided.

## Code availability

All code used for processing and statistically analysing data sets has been deposited in myoRDNA (https://github.com/MuscleOmicsLab/myoRDNA) for rDNA copy number analysis and the following GitHub repository (https://github.com/MuscleOmicsLab/WG-RRBS_pipeline_rDNA) for the bioinformatics pipelines.

